# *Mycobacterium tuberculosis* detection from oral swabs with Xpert MTB/RIF ULTRA: A pilot study

**DOI:** 10.1101/559815

**Authors:** Annelies Mesman, Roger Calderon, Martin Soto, Julia Coit, Juan Aliaga, Milagros Mendoza, Molly Franke

## Abstract

**Objective:** Diagnostic testing for tuberculosis depends on microbiological detection of *Mycobacterium tuberculosis* (*Mtb*) in sputum. For patients unable to expectorate sputum, such as children and individuals living with HIV, this poses barriers to rapid diagnosis and treatment initiation. Therefore, this study aimed to use oral swabs have as alternative sample type for *Mtb* detection via molecular testing.

**Results:** In a pilot study, we aimed to evaluate sensitivity of *Mtb* detection via oral swabs using Xpert MTB/RIF ULTRA. We enrolled 63 TB cases and controls from Lima, Peru, and detected *Mtb* from oral swabs with a sensitivity of 45% (95% confidence interval (CI): 29%-63%) and specificity of 100% (95% CI: 84%-100%) using liquid culture of sputum as reference test. Our current protocol will need optimization, but these results support future exploration of the use of oral swabs for *Mtb* detection.

## Introduction

Timely tuberculosis (TB) diagnosis and treatment initiation is hampered in patients unable to produce a sputum sample for microbiological confirmation of *Mycobacterium tuberculosis (Mtb) (e.g.*, children and individuals living with HIV). Recent research efforts have aimed to identify alternative, non-invasive sample types for molecular detection of TB disease in these patient groups.

The results of two recent studies (1,2) show successful molecular detection of *Mtb* from oral swabs, after collection in Tris-EDTA-Sucrose-SDS lysis buffer, followed by DNA extraction and IS6110 qPCR. Oral swabs would be an ideal alternative sample type to sputum -- collection is easy with a minimal risk on aerosol production. However, DNA extraction and in-house developed amplification assays are laborious procedures that require bio-safety level laboratory facilities. The rollout of Xpert MTB/RIF (Cepheid Sunnyvale, USA) and its subsequent recommended use by WHO as an initial diagnostic tool in 2013, introduced automated molecular testing to many laboratories in resource-limited settings. Recently, Cepheid introduced the novel MTB/RIF Ultra assay. This test is more sensitive than MTB/RIF, albeit at the cost of a slight reduction in specificity (3), and has improved *Mtb* detection in smear-negative respiratory samples (3,4), and therefore could be a valuable test for oral swab testing.

## Main text

### Study aim

In a small proof-of-concept pilot study we estimated the sensitivity and specificity of *Mtb* detection from oral swabs using Xpert MTB/RIF ULTRA (Cepheid, Sunnyvale, USA). A second aim of this pilot was to pilot three collection methods and storage procedures of swabs. In addition to the method described previously (1,2), which requires preparation of a buffer, we employed two other methods that could facilitate implementation of Xpert testing of oral swabs in low-resource settings.

### Methods

We recruited 33 adults diagnosed with pulmonary TB from TB clinics in Lima, Peru prior to anti-TB treatment initiation. Thirty controls (i.e., clinic attendees with no respiratory symptoms) were recruited from the same health centers. Participants were randomly assigned to one of three sample collection groups to assess collection methods and storage procedures: 1. Omniswab (Whatman, catalog #WB100035) in lysis buffer as described in (1,2); 2. Omniswab in phosphate buffered saline (PBS); 3. Sample capture on FTA cards (Whatman catalog #120237). We collected a single oral cheek swab from all TB patients and controls, and a sputum sample from TB patients. Sputum was used for microbiological confirmation via microscopy following Ziehl Neelsen staining and liquid culture (i.e., the gold reference standard; in BACTEC MGIT 960, BD Franklin Lakes, USA). For Xpert testing, 500-600 microliters of sample in PBS or lysis buffer was mixed in a 1:2 ratio with Xpert sample reagent buffer; sample reagent was directly added to FTA cards. Manufacturer’s guidelines were followed for all remaining steps of testing. Sensitivity and specificity and 95% confidence intervals (CI) were calculated via http://vassarstats.net.

### Results

All 33 TB patients had a positive sputum liquid culture result. The sputum of four patients tested negative in smear microscopy. The overall sensitivity for oral swab detection was 45% (95% confidence interval (CI): 29%-63%), with positive tests for 15/33 TB patients (Table 1). We observed a higher sensitivity for patients with a smear-positive sample compared to smear negative, although we included too few patients for statistical comparison. Sensitivity was similar across the three swab collection protocols, with swab collection in Tris-EDTA buffer resulting in *Mtb* detection in 6 of 10 patients, followed by FTA and PBS cards (5/13 and 4/10 respectively). Specificity was 100% for all groups (95% CI: 84.0%-100%), (26/26 of controls tested negative; 4 tests had invalid results due to power interruption).

**Table 1:** Xpert positivity per collection method for all TB cases and controls

|  | Lysis<br>buffer<br>n/N | FTA<br>n/N | PBS<br>n/N | Combined<br>n/N (%) | Sensitivity and<br>specificity<br>(95% CI) |
| --- | --- | --- | --- | --- | --- |
| SM<sup>a</sup> positive | 6/10 | 4/10 | 4/9 | 14/29 | 48% (30-67) |
| SM<sup>a</sup> negative | 0/0 | 1/3 | 0/1 | 1/4 | 25% (1.3-78) |
| All TB cases | 6/10 | 5/13 | 4/10 | 15/33 | 45% (29-63) |
| Control | 0/7 | 0/10 | 0/9 | 0/26 | 100% (84-100) |

<sup>a</sup> SM: smear

Three of the 33 patients were diagnosed with drug resistant TB based on culture-based drug sensitivity testing. We confirmed rifampin resistance with Xpert oral swab analysis for one of these patients, *Mtb* was not detected in the swabs of the other two patients.

### Discussion

These results support the possible utility of oral swab samples for TB detection. Luabeya et al reported that Omniswab tongue swabs yielded significant better results compared to cheek swabs, with a sensitivity of 71.2% for a single swab and 83.1% for two swabs in 59 patients (case definition based on Xpert or liquid culture) and 83.1% for single swab in another group of 89 patients (compared to Xpert MTB/RIF) (2). Their additional use of sputum Xpert as the reference test and tongue swab may in part explain the higher observed sensitivity in their study and would be interesting adjustments to the Xpert protocol. Their overall specificity was 91.5%, which would be below acceptable standards for a novel TB test, as described in the WHO target product profile (5). The authors hypothesize that the limited specificity was due to contamination errors in the PCR step. The automated integrated Xpert system is less prone to contamination, and the 100% specificity we measured in this pilot study is promising. Additionally, although current numbers are too small to draw any conclusions, these preliminary results suggest that other collection and storage procedures can be applied for oral swab testing. In conclusion, these data support the feasibility of *Mtb* detection in oral swabs using Xpert. Future work could explore the utility of oral swabs for the diagnosis of TB in patients unable to produce sputum and optimization of collection and storage strategies.

## Limitations

The major limitation of this study is the small sample size and pooling results of samples obtained by different collection methods. These numbers are insufficient to statistically test for differences in sensitivity across the collection methods. A second limitation is that we enrolled asymptomatic controls, whereas symptomatic individuals will be the most important group in which to evaluate the accuracy of this test. Finally, since we tested swabs from adult patients with culture-confirmed TB, this could overestimate the utility of this test for patients that would benefit most from oral swab testing (i.e., those with paucibacillary disease or who are unable to produce a sputum sample).

## Abbreviations

CI: confidence interval
MTB: Mycobacterium tuberculosis
PBS: phosphate buffered saline
SM: smear
TB: tuberculosis
RIF: rifampin
WHO: World Health Organization

## Declarations

### Availability of data and materials

Reported data are available on request from the corresponding author.

### Author’s contributions

AM: Study design, led the data analysis and interpretation and wrote the manuscript. RC: study design and data analysis. MS: study implementation and interpretation of results. JC: study implementation and interpretation of results JA: study implementation. MM: study implementation. MF: conceptualized and designed the study and contributed to writing and editing. All authors critically reviewed the manuscript and approved the final version.

### Ethics, consent and permissions

Study participants provided written informed consent to participate. Consent to publish is not applicable, we do not report individual patient data. This pilot was part of a study approved by Ethics Committee of Peru’s National Institute of Health and the Office of Human Research Administration at the Harvard Medical School.

## Acknowledgments

We are grateful for all participants in this study. We thank Socios En Salud for their contribution and dedication to health equity.

